# Amplification-free Library Preparation Improves Quality of Hi-C Analysis

**DOI:** 10.1101/562280

**Authors:** Longjian Niu, Wei Shen, Yingzhang Huang, Na He, Yuedong Zhang, Jialei Sun, Jing Wan, Daxin Jiang, Manyun Yang, Yu Chung Tse, Li Li, Chunhui Hou

## Abstract

PCR amplification of Hi-C libraries introduces unusable duplicates and results in a biased representation of chromatin interactions. We present a simplified, fast, and economically efficient Hi-C library preparation procedure that generates sufficient non-amplified ligation products for deep sequencing from 30 million *Drosophila* cells. Comprehensive analysis of the resulting data indicates that amplification-free Hi-C preserves higher complexity of chromatin interaction and lowers sequencing depth dramatically for the same number of unique paired reads. For human cells which has a large genome, this method recovers an amount of ligated fragments enough for direct high-throughput sequencing without amplification on as low as 250 thousand of cells. Comparison with published *in situ* Hi-C on millions of human cells reveals that amplification introduces distance-dependent amplification bias, which results in increasing background noise level against genomic distance. With amplification bias avoided, our method may produce a chromatin interaction network more faithfully reflecting the real three-dimensional genomic architecture.

## Introduction

Hi-C is a powerful tool for mapping interaction frequencies between chromatin fragments in a genome-wide and quantitative manner^1^. It compares the number of ligation events between each fragment pair in a large population of cells and thereby allows the identification of various genome structural features including compartments, topologically associating domains (TADs), and loops^1-5^. The main prerequisite for high-quality Hi-C analysis is the accurate quantification of chromatin interaction frequencies. The amount of DNA obtained in a typical Hi-C experiment is assumed to be insufficient for direct high-throughput sequencing. Thus, PCR amplification is a default step in Hi-C-related experiments^6^ to guarantee sequencing primer addition and to produce a sufficient amount of DNA for sequencing, especially for Hi-C experiments involving single or low cell numbers^7-18^.

The 3D nature of Hi-C deems sequencing depth and library complexity are two critical variables in evaluating the achievable resolution from Hi-C experiments, given a range of fragment sizes determined by the choice of restriction enzyme. Currently, many biological replicates and multiple rounds of PCR amplifications are required for high resolution genome architecture analysis in order to generate sufficient DNA with high enough complexity to represent the global chromatin interaction diversity within a cell population. Though universal primers are used, PCR amplification introduces duplicates and may skew Hi-C library composition, which may not be fully corrected by normalization methods^19-21^.

However, whether PCR amplification is really necessary and to what extent it changes the composition of Hi-C libraries has not been systematically examined. Here we present SAFE Hi-C, a *<u>s</u>*implified, *<u>a</u>*mplification-*f*ree, and economically *<u>e</u>*fficient process, in which paired reads generated by independent ligation events are saved. We tested this method on 30 million *Drosophila* S2 and 250 thousand human K562 cells. Comparison to traditional *in situ* Hi-C reveals amplification introduces distance-dependent bias in chromatin interaction frequency, which lowers the ratio between intra-TAD and inter-TAD interaction frequency especially for large genomes such as human cells. Efficient recovery of enough ligated fragments for direct high-throughput sequencing is important for accurate depicting the three dimensional genome architecture and the understanding of its functional role in transcription regulation, replication, genome stability and other critical biological activities related to chromatin.

## Results

### SAFE Hi-C Library Preparation and Sequencing

To determine how many biotin-labelled ligation events can be captured by streptavidin-conjugated beads, we carried out two SAFE Hi-C experiments as biological replicates on 30 million *Drosophila* S2 cells using DpnII (Fig. 1). We stripped off Hi-C ligation products from beads after addition of sequencing primers^22^ (Fig. 1b and Supplementary Table 1). Qubit quantification showed each SAFE Hi-C experiment recovered around 100 ng of ligated DNA, which roughly equals to 50 μl of 12 nM single strand DNA with a size around 500 bases, enough for at least 10 lanes of sequencing on the Illumina HiSeq X10 platform. We also conducted traditional *in situ* Hi-C experiments in replicates for comparison and amplified Hi-C ligation products from diluted beads using different numbers of PCR amplification cycles (4, 8, 12, 16 and 20) to produce similar amounts of DNA as in the SAFE Hi-C experiments (Supplementary Table 1).

**Fig. 1.**
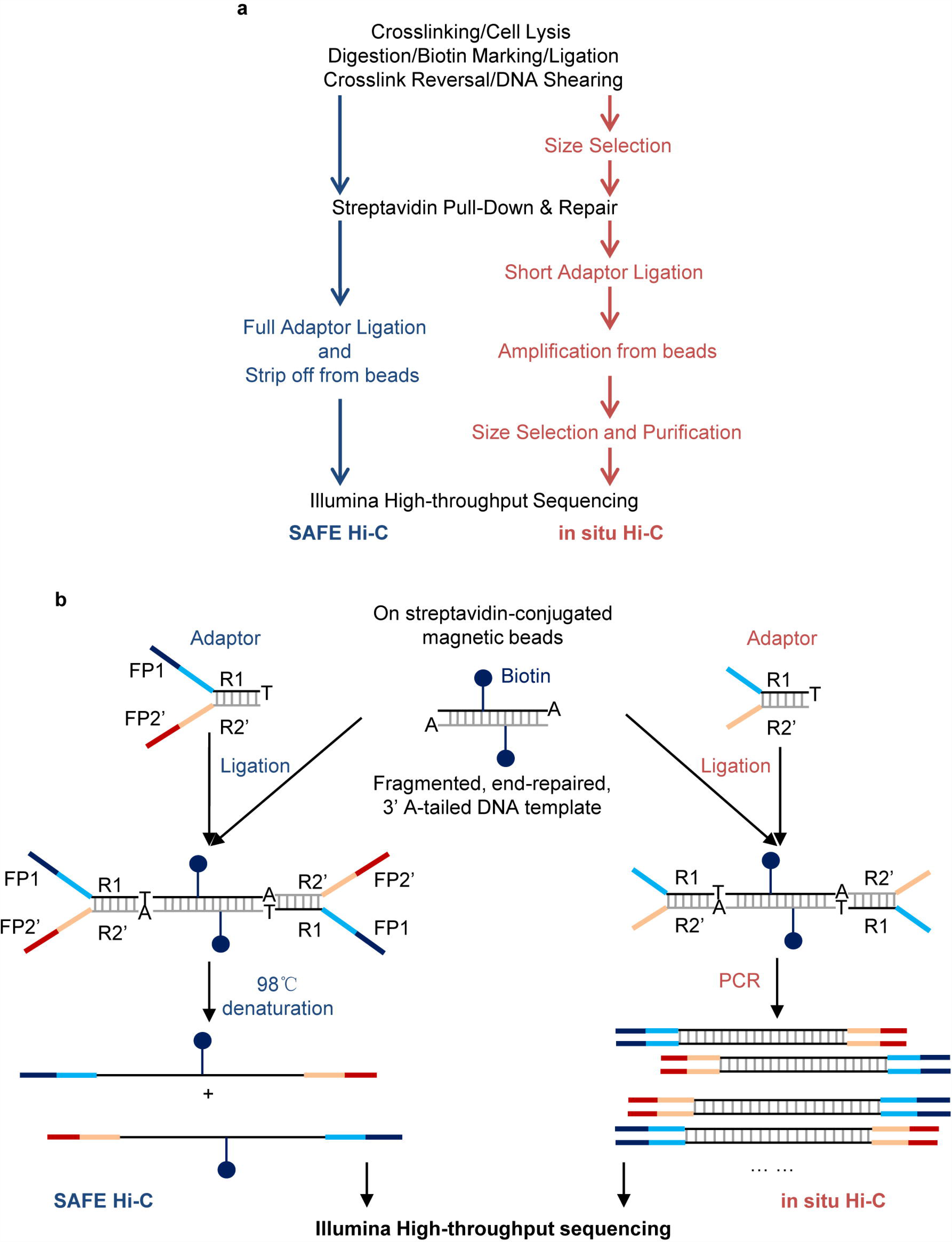
Procedures of SAFE Hi-C and traditional *in situ* Hi-C. **a** Side-by-side comparison of SAFE Hi-C and *in situ* Hi-C procedures. The black text shows shared steps in both methods, blue and red texts correspond steps specific for SAFE Hi-C and *in situ* Hi-C, respectively. **b** In the preparation of both *in situ* Hi-C and SAFE Hi-C libraries, partially complementary adapters with a 3’ thymine (T) overhang are ligated onto repaired and 3’ adenine (A)-tailed DNA captured on streptavidin beads. The sequence of adaptors is listed in Supplementary Table 1.

All libraries were sequenced on an Illumina HiSeq X10 instrument and aligned to the reference genome using bowtie 2.0. The length of sequenced fragments in all Hi-C libraries ranged from 200-750 bp, and peaked around 370 bp (Supplementary Fig. 1), consistent with the fact that the median length of DpnII fragments is 194 bp. Global chromatin interaction frequencies were highly correlated between biological replicates and between different pairs of libraries, with the lowest Pearson’s correlation value of 0.994 (Supplementary Fig. 2). We combined biological replicates and obtained 338, 246, 220, 232, 238 and 248 million aligned paired reads (Supplementary Table 2). Datasets were normalized as described^5,23^ for further analysis. The ratio of cis-and trans-unique paired reads, generally considered as a proxy indicator of the quality of a Hi-C library, was 13.5 for SAFE Hi-C and approximately the same for amplified libraries (Fig. 2a). After PCR amplification, fragment pairs with lower (<∼ 42%) or higher (>∼ 42%) GC content were significantly under- or over-represented compared to SAFE Hi-C, respectively (p<10^-40^, Mann–Whitney *U* test, Supplementary Fig. 3).

**Fig. 2.**
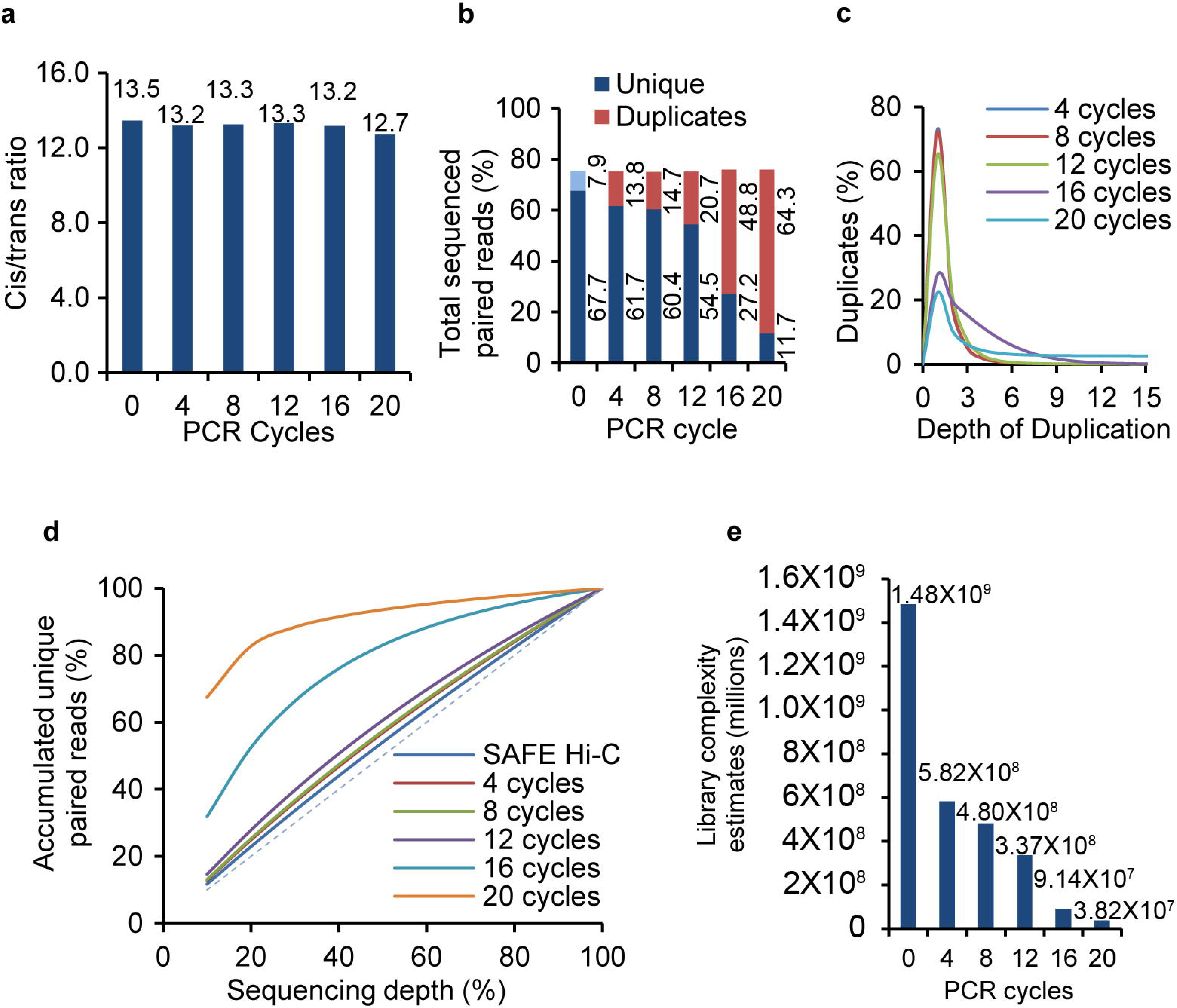
Amplification increases PCR duplicates and reduces Hi-C library complexity of the *Drosophila* genome. **a** Cis paired reads are uniquely mapped on the same chromosome, and trans paired reads are mapped on different chromosomes. SAFE Hi-C is referred to as 0 PCR cycle in figures. **b** Percentage of unique paired reads and duplicates that can be aligned out of the total sequenced paired reads. The light blue bar shows the PCR duplicates in SAFE Hi-C libraries, which we kept as unique for no amplification was involved in the library preparation process. **c** Frequencies of duplicate ligates introduced by PCR amplification. **d** Accumulated percentage of unique paired reads against the percentage of sequencing depth. **e** Library complexity estimates. For SAFE Hi-C, the complexity was estimated as described in Methods.

**Fig. 3.**
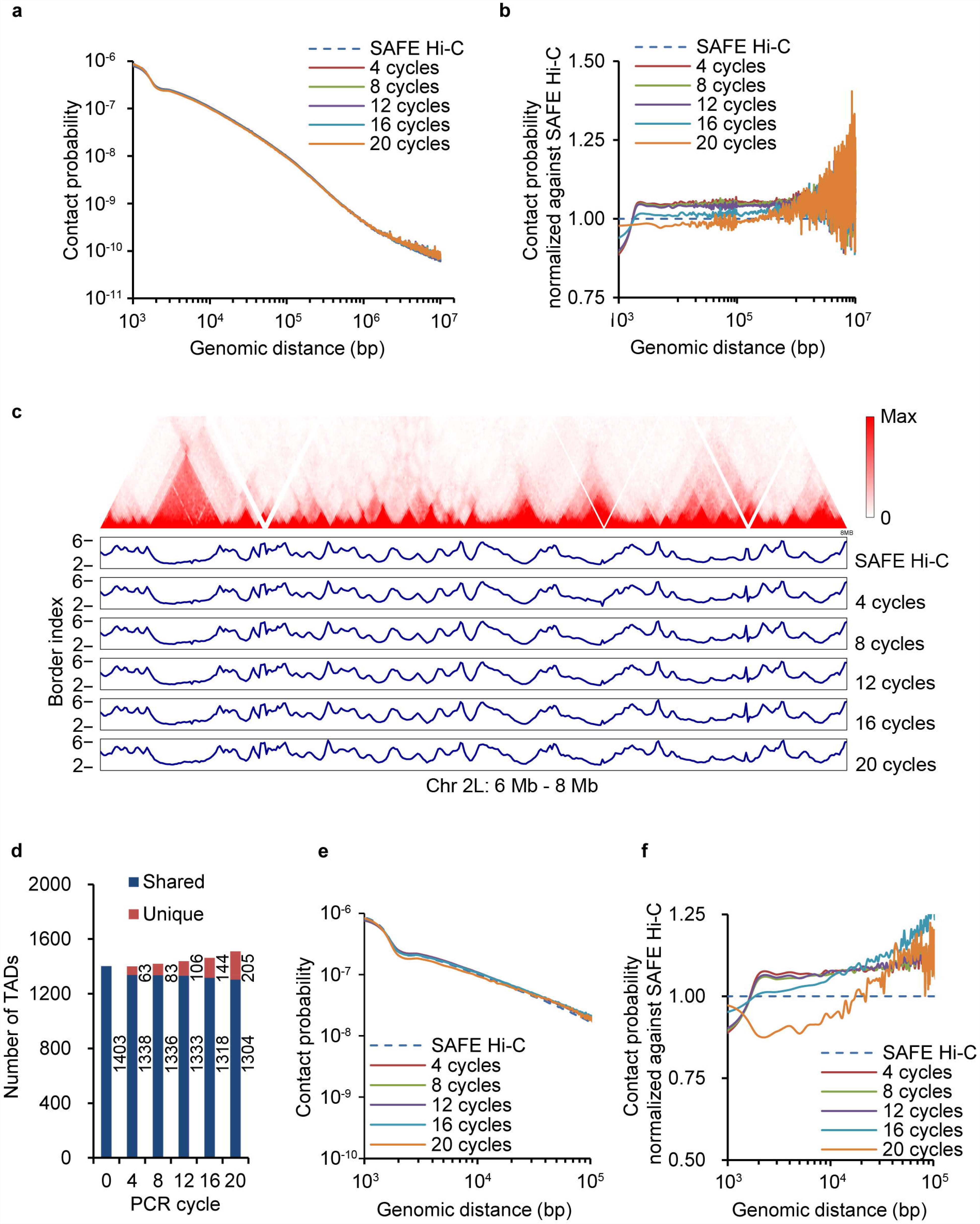
Distance-related amplification bias and TADs identification in *Drosophila* genome. **a** Chromatin interaction frequency as a function of genomic distance averaged across the genome for the *Drosophila* genome. **b** Average chromatin interaction frequency across the genome normalized against SAFE Hi-C for *Drosophila* genome. **c** Hi-C interaction heatmap for the region of chromosome 2L from 6 Mb to 8 Mb is shown. Border index values for SAFE Hi-C and amplified Hi-C libraries are compared. **d** Dark blue bars show the number of TADs shared with SAFE Hi-C, red bars show the number of TADs uniquely found for amplified Hi-C libraries. **e** Chromatin interaction frequency as a function of genomic distance averaged within the TADs. **f** Average chromatin interaction frequency within the TADs normalized against SAFE Hi-C.

After successfully applying SAFE Hi-C on 30 million *Drosophila* cells whose chromatin content is roughly equal to 1 million human cells, next we determined the lowest number of human cells needed for SAFE Hi-C. 250 thousand and 100 thousand human K562 cells were tested. SAFE Hi-C worked only with 250 thousand cells from which we recovered 15μl of 4 nM single strand DNA with a size around 575 bases. This is about 1/10 of the DNA recovered from 30 million *Drosophila* S2 cells. We think this may be due to the higher chance of DNA loss frequently experienced with experiment using small amount of starting material. One quarter of a lane on the Illumina HiSeq X10 was used for sequencing the human K562 SAFE Hi-C library generating 121 million of paired reads, out of which 106 million (87.4%) aligned successfully to the human reference genome (Supplementary Table 3). A similar number of chromatin interactions from *in situ* Hi-C on K562 cells previously published by Rao et al. was downloaded and used for comparison.

### SAFE Hi-C Avoids Removal of PCR Duplicates

PCR amplification introduces extra duplicates to the Hi-C library, which lowers the percentage of unique paired reads. For SAFE Hi-C, we define “suspected PCR duplicates” as those sharing identical sequences at both ends. These duplicates are most likely derived from ligation events of fragment pairs with same sequences in millions of cells, which could happen at a high rate especially when the size of most fragments is short. PCR duplicates can also be generated from DNA cluster generation on the Illumina sequencing machine and are considered as “optical duplicates”. This accounts for less than 1% of the total sequenced paired reads (Supplementary Table 2 and 3) and were excluded in further analysis.

About 8% of total mapped paired reads were “suspected PCR duplicates” for *Drosophila* S2 SAFE Hi-C libraries (Fig. 2b, light blue bar) and were kept for Hi-C analysis because no amplification was involved. However, for amplified libraries, duplicates from independent ligations cannot be distinguished from those introduced by PCR amplification, thus all were considered arising from single ligation events. As expected, the proportion of PCR duplicates positively correlates with the number of amplification cycles (Fig. 2b, red bar). The percentage of duplicates increased to 14%, 15%, 21%, 49% and 64% after 4, 8, 12, 16 and 20 cycles of amplification, respectively (Fig. 2b, red bar). Correspondingly, the percentage of non-duplicate paired reads decreased dramatically as amplification cycles increased, especially after 16 and 20 cycles (Fig. 2b, dark blue bar). We also calculated duplicates and valid paired read percentages for all mappable paired reads (Supplementary Fig. 4). PCR duplicate depth analysis shows that most duplicates have two copies for all libraries (Fig. 2c), and the percentage of amplified ligation products of higher duplicate depth increased considerably after 16 and 20 cycles of amplification (Fig. 2c).

**Fig. 4.**
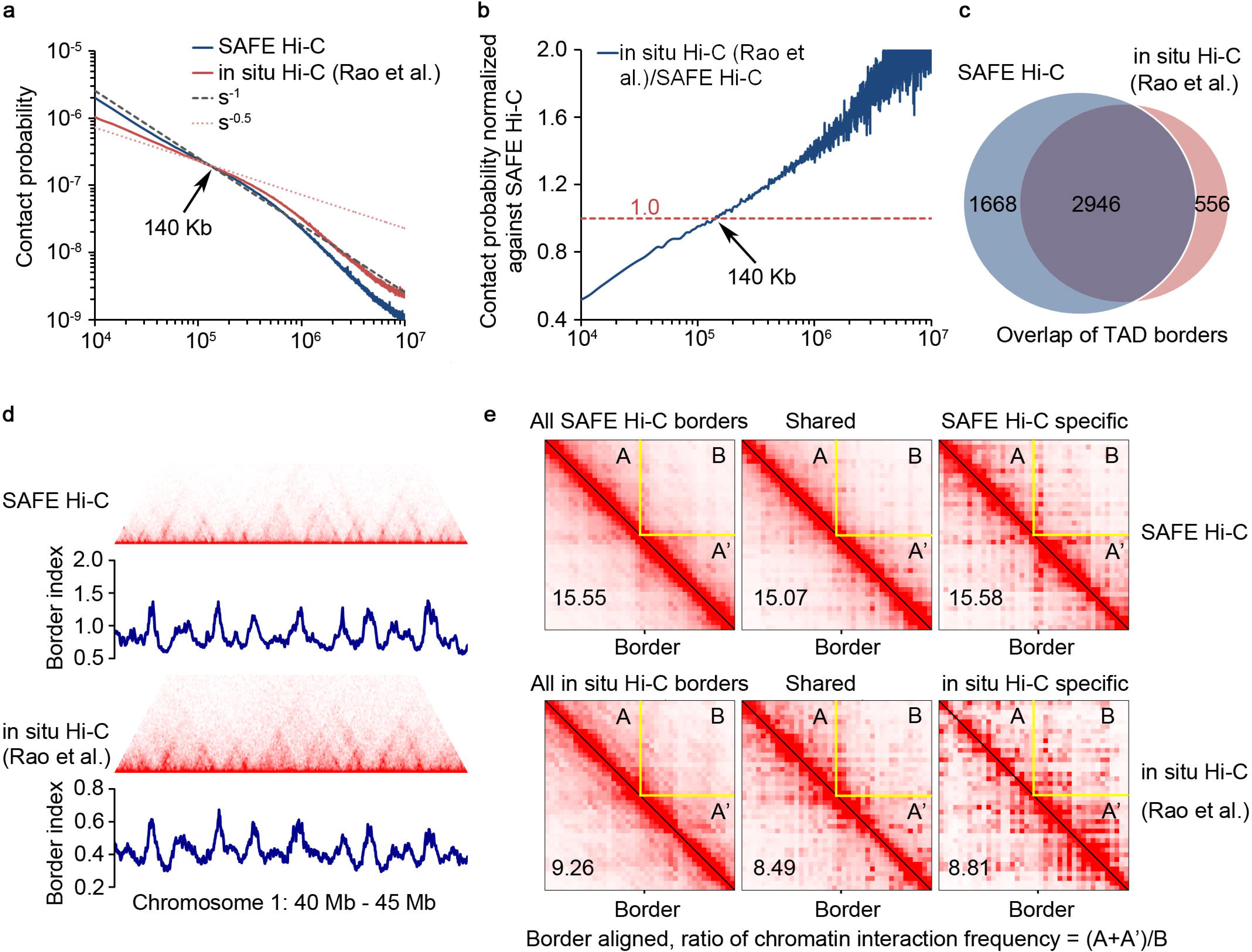
SAFE Hi-C on 250 thousand human K562 cells. **a** Comparison of chromatin interaction frequency against the genomic distance between SAFE Hi-C (blue line) and *in situ* Hi-C (red line). s^-1^ (black dashed line) and s^-0.5^ (red dotted line) represent the predicted fractal globule and mitotic states, respectively. The turning point of chromatin interactions on the decaying curves for SAFE Hi-C and *in situ* Hi-C is indicated by an arrowhead. **b** Average chromatin interaction frequency of *in situ* Hi-C normalized against SAFE Hi-C across the genome. The crossing point is shown by an arrowhead. **c** Venn diagram shows the overlap between TADs identified for SAFE Hi-C and *in situ* Hi-C. **d** Hi-C interaction heatmap and border index comparison between SAFE Hi-C and *in situ* Hi-C for a region of chromosome 1 from 40 Mb to 45 Mb. **e** Borders are aligned for all, shared and specific TADs. The chromatin contact frequency in flanking TADs (A+A’) is divided by that of the inter-TADs (B), the ratios of (A+A’)/B for aggregated borders are shown at the bottom left in each heatmap.

### SAFE Hi-C Increases Library Complexity

For *Drosophila* S2 SAFE Hi-C libraries, the percentage of unique pair reads correlates almost linearly with sequencing depth (Fig. 2d), suggesting that the library complexity was far from exhausted at current sequencing depth. After amplification, PCR duplicates cannot be kept and their percentage increases dramatically as the cycle of amplification increases (Fig. 2b). Consistently, the estimated library complexity dramatically dropped from 1.5 billion for SAFE Hi-C libraries to 0.58 billion for *in situ* Hi-C libraries after just 4 cycles of amplification (Fig. 2e).

### Amplification Bias Is Genomic Distance-Dependent

Chromatin interaction frequency is inversely correlated with genomic distance. Amplification resulted in a moderate change on the decaying pattern of chromatin interactions at any genomic distance for traditional *in situ* Hi-Cs on *Drosophila* S2 (Fig. 3a, b). However, for K562, the changes were obvious (4a, b). After normalization against SAFE Hi-C, we found that the relative chromatin interaction frequency starts lower (∼ 0.9) at 1kb and becomes higher beyond 3 kb for most amplified libraries in *Drosophila* S2 cells, except for amplification of 20 cycles (Fig. 3b), suggesting short distance ligations are generally underrepresented after amplification even after limited number of amplification cycles.

In comparison to *Drosophila S2*, amplification introduced more obvious biases for the human genome. Compared to published *in situ* Hi-C on human K562 cells, chromatin interaction frequency from SAFE Hi-C decays at a rate much closer to the predicted fractal globular model (s^-1^, Fig. 4a), while *in situ* Hi-C decays at a rate closer to s^-0.5^ within the genomic distance of 1 Mb (Fig. 4a). When normalized against SAFE Hi-C, the relative chromatin interaction frequency from *in situ* Hi-C starts even lower (∼ 0.5) at 10kb and continuously increases and crosses with SAFE Hi-C at a genomic distance of 140kb (Fig. 4b).

This comparison reveals an unexpected bias in Hi-C library amplification, which could be due to the competition of complementary fragment hybridization and primer annealing. It is reasonable to expect that pairs of ligated fragments of high concentration tend to hybridize to themselves instead of primers. The increasing bias over distance also suggests that background noise could be more significant over a longer distance.

### SAFE Hi-C Maintains a Higher Contrast Ratio of the Intra-TAD and Inter-TADs Chromatin Interaction Frequency in the Human Genome

So far, all characterizations of TADs, compartments and loops have been based on amplified Hi-Cs. However, the amplification effects on such analysis have not been evaluated experimentally. With the development of SAFE Hi-C, we are able to show if and to what extent amplification affects TADs, compartments and loops analysis.

We first plotted heatmaps for *Drosophila* S2 cells with normalized datasets and identified TADs at 5kb resolution (Fig. 3c and Supplementary Fig. 5). Overall, we observed minimal variation in border strength across the genome after amplification (Fig. 3c). Consistently, the number of identified TADs was similar and most of them were conserved (Fig. 3d). These results together suggest at least SAFE Hi-C is as reliable as traditional in situ Hi-C for TAD characterization.

We further calculated the interaction frequency vs distance within TADs (Fig. 3e). Similar to a global decaying pattern, normalization against SAFE Hi-C within TADs showed that chromatin interaction frequency was underrepresented within and overrepresented beyond 3 kb for most amplified libraries (Fig. 3f). Recently, sub-kb resolution Hi-C identified 4,123 TADs in the *Drosophila* genome, with TADs as small as 3 kb^24^. Our SAFE Hi-C results also indicate that high frequency of interactions in the 3 kb range may be an important feature of the *Drosophila* genome.

Next, we identified TADs for SAFE Hi-C and *in situ* Hi-C on human K562 cells. Most of the TADs identified overlapped (2946), however, we found many more specific TADs for SAFE Hi-C (1668) as compared with *in situ* Hi-C (556) (Fig. 4c). Visual inspection shows a very similar pattern of border index for both experiments (Fig. 4d). However, for *in situ* Hi-C border index peaks are frequently less sharp (arrowhead, Fig. 4d) and the shape of possible TADs is generally less clean compared with SAFE Hi-C (heatmap, Fig. 4d). The border index values are lower for *in situ* Hi-C as well (Fig. 4d). We aligned the borders of all identified TADs, shared TADs and specific TADs for SAFE Hi-C and *in situ* Hi-C and calculated the ratio of chromatin interaction density of intra-TAD over inter-TAD (Fig. 4e). The ratios of SAFE Hi-C (15.55, 15.07 and 15.58) are consistently higher than the values for *in situ* Hi-C (9.26, 8.49 and 8.81) (bottom left, Fig. 4e). These results together show that amplification weakens the contrast between TADs and inter-TAD regions, which could be caused by biased amplification over genomic distance as shown in Fig. 4a and 4b, and thus underlines the importance of avoiding amplification in order to further improve the quality of analysis.

### Amplification Effects on Compartment Analysis and Chromatin Loop Identification

We characterized compartments for each chromosome arm of the *Drosophila* genome (Supplementary Fig. 6). The span of absolute eigenvalues decreased for amplified Hi-C (Supplementary Fig. 7), suggesting SAFE Hi-C may somehow improve compartments analysis. For human K562 cells, the eigenvalues correlated highly for SAFE Hi-C and *in situ* Hi-C (Supplementary Fig. 8). Together, these results show that the effect of amplification on compartment analysis is less obvious than for TADs identification.

Finally, we identified long-range chromatin interactions at 5 kb resolution for the *Drosophila* genome (q value < 0.1) but not for human genome because the resolution is too low for meaningful and reliable chromatin loop identification. Signal-to-noise ratio was calculated as described^5^. The number of identified loops negatively correlated with the number of amplification cycles (Supplementary Fig. 9 and 10a). Approximately 43% of loops from each amplified Hi-C library (Supplementary Fig. 10a, dark blue bar) overlapped with those identified using SAFE Hi-C. The aggregate P2LL values and ZscoreLL for all loops identified for both SAFE Hi-C and amplified libraries changed little (Supplementary Fig. 10b, c). For shared loops, the P2LL values rose as the number of PCR cycles increased (Supplementary Fig. 10d, upper panel), suggesting amplification favored these interactions. However, for loops lost after amplification, the P2LL values decreased (Supplementary Fig. 10d, lower panel), suggesting under-amplification of these interactions.

## Discussion

In addition to the introduction of PCR duplicates and a significant reduction in library complexity, our results show that amplification has a different effect on the three-dimensional genome architecture analysis for a small (*Drosophila*) and big (human) genomes. For small genomes like *Drosophila*, amplification affects the characterization of loops and to lesser extent compartments, but shows little effect on accurate TAD identification when the same number of raw sequencing paired reads were used.

Though SAFE Hi-C was initially tested on *Drosophila* genome which is much smaller than typical mammalian genomes, it is also suitable for large genomes due to its ability to maintain the original complexity and integrity of chromatin interaction diversity, effectively lowering sequencing depth and saving labor and cost. We estimate that 1-2 million mammalian cells digested with DpnII will generate the same amount of Hi-C ligation products as 30 million *Drosophila* cells, which will be enough for at least ten lanes of sequencing on the Illumina X10 platform. Theoretically, the complexity of a single SAFE Hi-C library will be in the hundreds of billions if the number of single molecular DNA ligation products was calculated in a library made from 1-2 million mouse or human cells.

We tested the low limit of human cell number for SAFE Hi-C. With 250 thousand of K562 cells, we successfully recovered ligates enough for at least one lane of sequencing on the Illumina X10 platform. With a lower number of cells, the chance of DNA loss increased during library preparation. Our data analysis reveals that avoiding amplification significantly improves TAD identification for the human genome. It is reasonable to conclude that the genomic distance dependent bias introduced by amplification may very likely compromise the quality of TAD analysis.

In sum, by avoiding amplification, SAFE Hi-C can be used to improve the quality of Hi-C analysis as well as to save time, reagents and reduce cost. In cases when the available cell number is too low, SAFE Hi-C is also compatible with PCR amplification to ensure enough sequencing material can be generated. Finally, other Hi-C methods using enzymes like DNase or MNase may also be simplified and be amplification-free.

## Methods

### Cell culture

S2 cells were cultured in Schneider’s Medium (Gibco, 21720024) supplemented with 10% heat inactivated FBS (Sigma, F7524) and 1% Penicillin/Streptomycin P/S (Sigma, P0781) at 27°C.

K562 cells were incubated in 1× RPMI1640 media supplemented with 10% FBS at 37°C with 5% CO2.

### *In situ* Hi-C

*In situ* Hi-C was carried out as described. Cells were crosslinked with 1% formaldehyde then lysed to collect nuclei. Pelleted nuclei were digested with DpnII restriction enzyme (NEB, R0147). The restriction fragment overhangs were filled and marked with biotin-labelled dATP (Thermo Fisher, 19524016) and dCTP, dTTP and dGTP before ligation. DNA was reverse crosslinked, purified and fragmented by sonication on a Covaris sonicator. Biotin labelled DNA was pulled down on Streptavidin Dynabeads (NEB, S1420S). After DNA repair and 3’ A addition, SHORT Y-Adaptor (Supplementary Table 1) was added. Diluted DNA on Dynabeads was used for PCR amplification (4, 8, 12,16 and 20 cycles) to produce similar amounts of DNA for sequencing on the Illumina HiSeq X10 platform (PE 2×150 bp reads).

### SAFE Hi-C

SAFE Hi-C is a modification of in situ Hi-C^5,25^. Cells were crosslinked with 1% formaldehyde for 10 min at room temperature (RT). The reaction was stopped by adding 1/10 volume of 2.5 M glycine. Up to 30 million crosslinked cells were resuspended in 500 µL of ice-cold Hi-C lysis buffer and rotated at 4 °C for 30 min. Nuclei were pelleted at 4 °C for 5 min at 2,500 relative centrifugal force (RCF), and the supernatant was discarded. Pelleted nuclei were washed once with 500 µL of ice-cold Hi-C lysis buffer. The supernatant was removed again, and the pellet was resuspended in 100 µL of 0.5% SDS and incubated at 62°C for 10 min with no shaking or rotation. 285 µL of water and 50 µL of 10% Triton X-100 were added, and samples were rotated at 37°C for 15 min to quench the SDS. 50 µL of NEB Buffer 3.1 and 20 µL of 10 U/µL DpnII restriction enzyme (NEB, R0147) were then added, and the sample was rotated at 37°C for 4 h. DpnII was then heat inactivated at 62°C for 20 min with no shaking or rotation. To fill-in the restriction fragment overhangs and mark the DNA ends with biotin, 52 µL of incorporation master mix was then added: 37.5 µL of 0.4 mM biotin–dATP (Thermo Fisher, 19524016); 4.5 µL of dCTP, dGTP, and dTTP mix at 10 mM each; and 10 µL of 5 U/µL DNA Polymerase I Large (Klenow) Fragment (NEB, M0210). The reactions were then rotated at 37°C for 45 min. 948 µL of ligation master mix was then added: 150 µL of 10× NEB T4 DNA ligase buffer with 10 mM ATP (NEB, B0202), 125 µL of 10% Triton X-100, 3 µL of 50 mg/mL BSA (Thermo Fisher, AM2616), 10 µL of 400 U/µL T4 DNA Ligase (NEB, M0202), and 660 µL of water. The reactions were then rotated at 16 °C for 4 h and room temperature for 1 h. 45 µl of 10% SDS and 55 µl of 20 mg/ml Proteinase K were added for crosslinking reversal. Incubate at 55°C for at least 2 hours (overnight recommended). DNA was purified by phenol:chloroform:isoamyl alcohol (25:24:1) extraction. Purified DNA in solution was transferred into a 1.5 ml tube and sonicated to 400 bp on a Covaris sonicator. Biotin labelled DNA was pulled-down on Streptavidin Dynabeads (NEB, S1420S). After DNA repair and 3’ A addition, Full Y-Adaptor (Supplementary Table 1) was added. DNA -on Dynabeads was resuspended in 100 μl of 0.8× PCR Buffer and incubated at 98°C for 10 min before being cooled off in ice water. The supernatant was recovered, quantified, and used for direct sequencing on the Illumina HiSeq X10 platform (PE 2×150 bp reads). SAFE Hi-C on human K562 cells was carried out similarly with the reagents reduced in proportion to the estimated chromatin contents, not the cell number.

### Data processing

We chose dm3 and hg19 as the reference genomes to align sequenced reads. Mapping, filtration, duplication removal, construction and normalization of contact matrices and basic library statistics of reads from all experiments were processed using the Juicer pipeline^26^. For SAFE Hi-C, the analysis should only remove optical duplicates, which is caused by sequencing when a single cluster of reads is part of two adjacent tiles’ on the same slide and used to compute two read calls separately. We used a modified AWK script derived from juicer’s dups.awk script which removes duplicates and judges the source of duplicates (PCR or optical) to remove only optical duplicates. We set 1 as map quality threshold and all downstream analyses were based on KR normalized matrices ^23^ which ensures that each row and column of the contact matrix sums to the same value.

### Analysis of PCR duplication rates for *Drosophila* S2 libraries

For duplication depth analysis, we used a modified AWK script derived from juicer’s dups.awk script to count the duplicate number of each duplicated contact. We also tested different wobble number (0, 1, 2, and 3) to process the deduplication step for each library. We show that SAFE Hi-C libraries have high library complexity in standard deduplication (wobble=4) process and even higher when set wobble to 0.

### Library complexity estimation for *Drosophila* S2 libraries

Estimation of library complexity has been described before^5,27^. For SAFE Hi-C libraries, we computed the “PCR duplication rate” (although this library does not contain real PCR duplicates) to estimate library complexity.

### Topologically associating domain identification

Identification of TADs in *Drosophila* S2 cells was processed by Juicer Arrowhead algorithm at 5kb resolution. For the identification of TADs in human K562 cells, border strength index was calculated at 25kb resolution with a moving block size of 8 bins. TAD borders of human K562 dataset were defined by calling peaks through R package pracma. We used bedtools^28^ intersect command to call overlapped TADs and the overlapped region between two overlapped TADs should span at least 90 percent of each TAD range (command: bedtools intersect -f 0.9 -F 0.9 -sortout -a $tad_1 -b $tad_2)

### Compartment analysis

Method of compartment analysis has been described before^6^. We used the Pearson’s and eigenvector command of Juicer tools to obtain the Pearson’s correlation matrix and eigenvector at 10 kb resolution.

### Long-range chromatin interaction calling and aggregate peak analysis

Loops were identified at 5 kb and 10 kb resolution respectively using Juicer’s HiCCUPs algorithm^26^ (parameters: -m 2048 -r 5000,10000 -k KR--ignore_sparsity). P2LL (Peak to Lower Left) and ZscoreLL (Zscore Lower Left) are defined to measure the enrichment of HiCCUPs peaks during aggregate peak analysis (APA). P2LL is the ratio of the central pixel to the mean of the pixels in the lower left corner. ZscoreLL stands for the Z-score of the central pixel relative to all of the pixels in the lower left corner. Note that, for P2LL scatter plot (fig*), the P2LL value is the ratio of each peaks’ pixel to the expect value of lower left.

## Data availability

*Drosophila* S2 raw Illumina reads are available at

https://www.ncbi.nlm.nih.gov/bioproject/PRJNA470784.

Accession code: PRJNA470784

Human K562 raw Illumina reads are available at

https://www.ncbi.nlm.nih.gov/bioproject/524051

Accession code: PRJNA524051

## Code availability

All custom codes are available at

https://github.com/shenscore/Safe_Hi-C_script.

## Acknowledgments

We gratefully acknowledge financial support from the National Key R&D Program of China (2018YFC1004500), National Natural Science Foundation of China (31571347 to C.H., 31771430 to L. L. and 31671409 to Y.T.), Guangdong Science and Technology Department (2016A030313642 to C.H.), Shenzhen Science and Technology Innovation Commission (JCYJ20150529152146478 to C.H., JCYJ20170307105005654 to Y.T.), Huazhong Agricultural University Scientific & Technological Self-innovation Foundation (to L. L.) and the Thousand Talent Youth Program (to C.H.). We thank Dr. Victor G. Corces for helpful advices on manuscript preparation and Dr. Edwin Cheung for text editing.

## Author contributions

C.H. and L.N. conceived the study and designed the experiments; L.N. performed the experiments with help from Y.Z., Y.H., J.S., D.J., M. Y. and Y.T.; W.S., N.H. and J. W. carried out the data analysis; L. L. supervised the data analysis; C.H. supervised the study; C.H. wrote the manuscript with input from all authors.

## Legends for Supplementary Figures and Tables

**Supplementary Fig. 1** Size distribution of ligation products from Hi-Cs on *Drosophila* S2. Size was calculated based on the distance of each end of a paired read to its closest DpnII sequence at the genomic site.

**Supplementary Fig. 2** Pearson correlation between biological duplicates and between SAFE Hi-C and *in situ* Hi-C libraries with the different number of amplification cycles for *Drosophila* S2.

**Supplementary Fig. 3** Amplification effect on paired reads of different GC content. Paired reads with GC content from 20%-80% are equally divided into 100 equal bins. (**a**) Ratio of paired reads as a function of GC content for SAFE Hi-C and *in situ* Hi-C libraries with the different number of amplification cycles. Ratio is the number of paired reads of each specific bin over the number of total paired reads. (**b**) Ratio of paired reads was normalized against SAFE Hi-C.

**Supplementary Fig. 4** Percentage of unique paired reads and duplicates of the total alignable paired reads. The light blue bar shows the “suspected PCR duplicates” that were kept as unique.

**Supplementary Fig. 5** 5 kb heatmaps for each chromosome arm of *Drosophila* S2 genome.

**Supplementary Fig. 6** 10 kb resolution compartments for each chromosome arm of *Drosophila* S2 genome.

**Supplementary Fig. 7** Pearson correlation analysis of compartment eigenvalues from SAFE Hi-C and amplified Hi-Cs for *Drosophila* S2 cells.

**Supplementary Fig. 8** Pearson correlation analysis of compartment eigenvalues of SAFE Hi-C and *in situ* Hi-C on human K562 cells.

**Supplementary Fig. 9** Examples of long-range chromatin interaction loops identified on Chr 2L between 10.2 – 11.0 Mb for the *Drosophila* genome by SAFE Hi-C and amplified Hi-Cs. Loops identified for SAFE Hi-C and for amplified Hi-Cs are marked by blue and green circles, respectively. Short vertical lines above heatmaps indicate CTCF binding sites. *Drosophila* S2 CTCF binding sites are downloaded from modENCODE.

**Supplementary Fig. 10** Comparison of long-range chromatin interactions loops identified for the *Drosophila* genome by SAFE Hi-C and amplified libraries. (**a**) Number of long-range chromatin interaction loops. Dark blue bars show the number of loops shared with SAFE Hi-C, red bars show the number of loops uniquely found for amplified Hi-C libraries. (**b**) Average signal-to-noise values. P2LL is calculated as described in Methods. (**c**) Average ZscoreLL for identified chromatin loops. ZscoreLL is calculated as described in Methods. (**d**) Correlation analysis of P2LL values for loops shared with SAFE Hi-C (upper panel) and for loops found for SAFE Hi-C but lost after amplification (lower panel). Red dots show loops with higher P2LL value and blue ones with lower values after amplification compared to SAFE Hi-C.

**Supplementary Table 1** List of adaptors and primer sequences.

**Supplementary Table 2** Libraries sequencing statistics for *Drosophila* S2 cells.

**Supplementary Table 3** SAFE Hi-C and *in situ* Hi-C sequencing statistics for human K562 cells.

