## Supplementary figures and images for "Amplification-free Library Preparation Improves Quality of Hi-C Analysis"

### Supplementary Figure 1

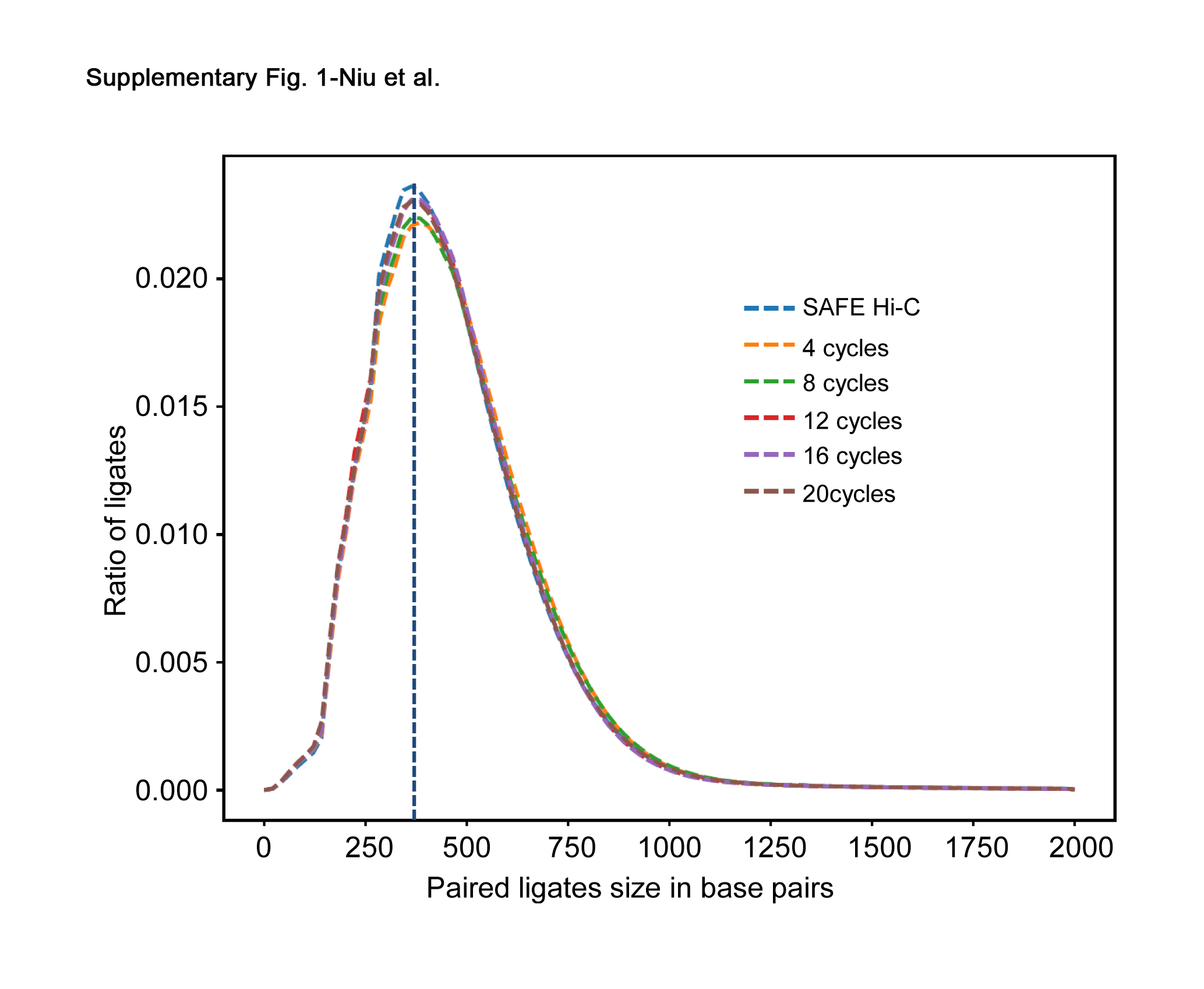

### Supplementary Figure 2

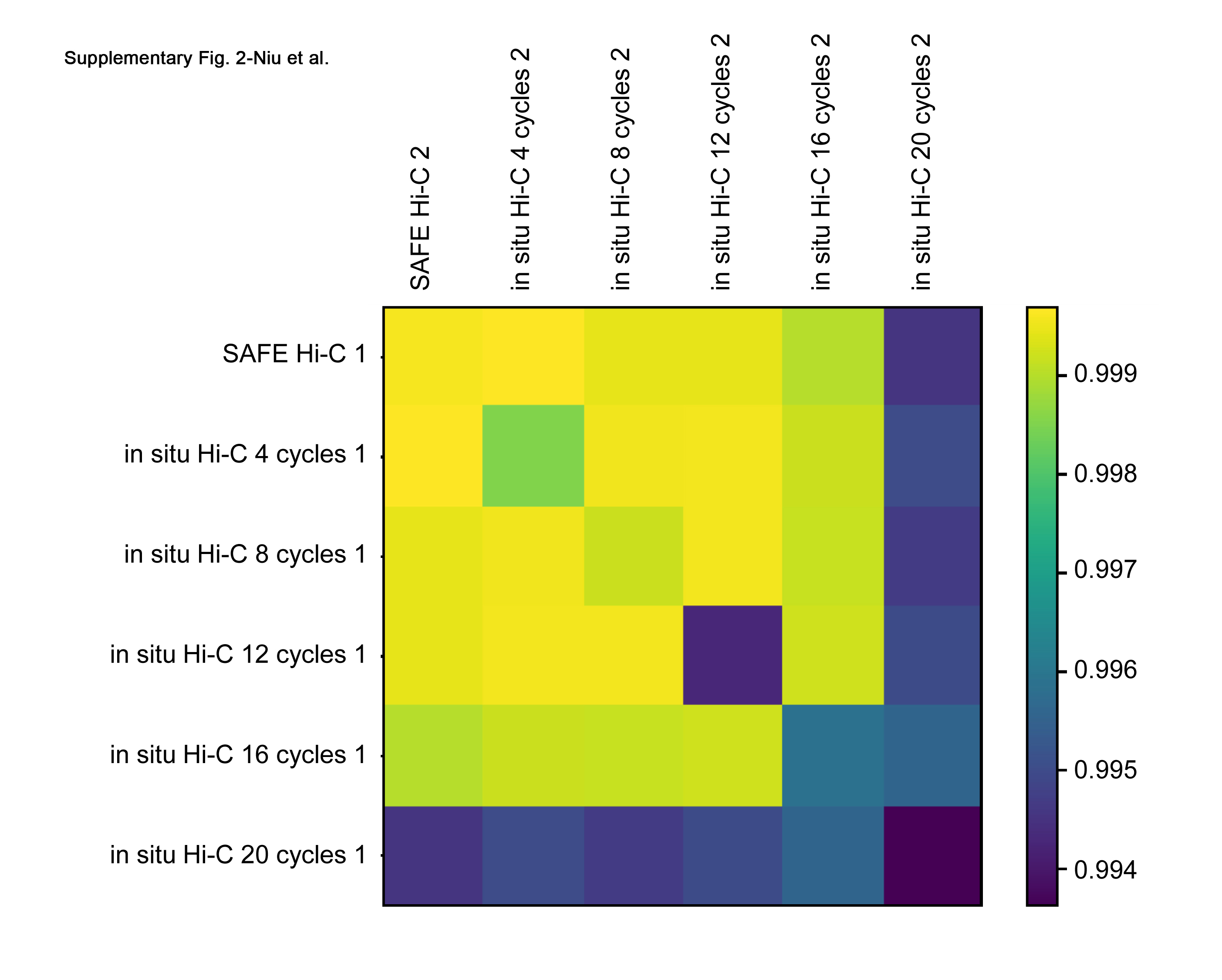

### Supplementary Figure 3

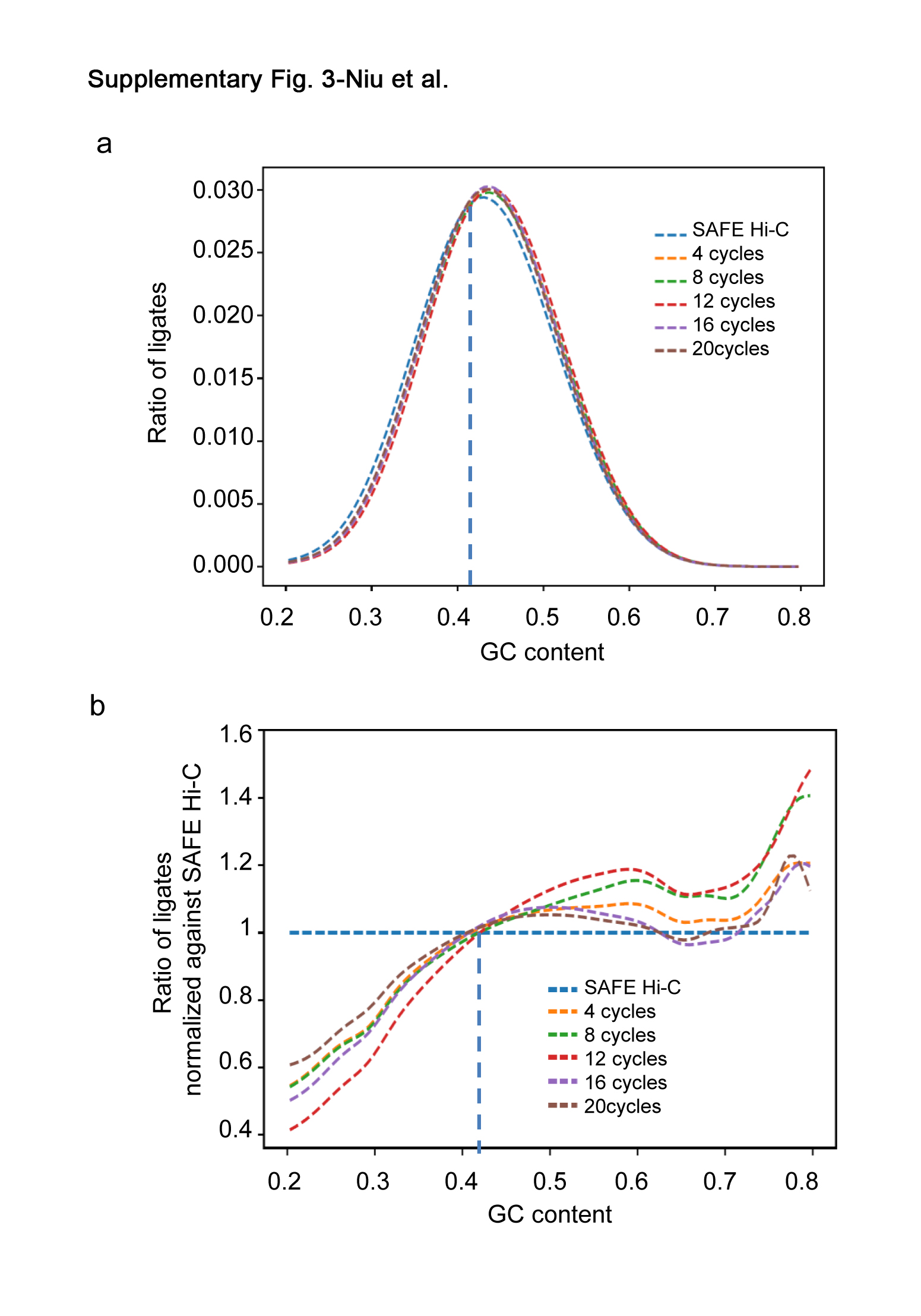

### Supplementary Figure 4

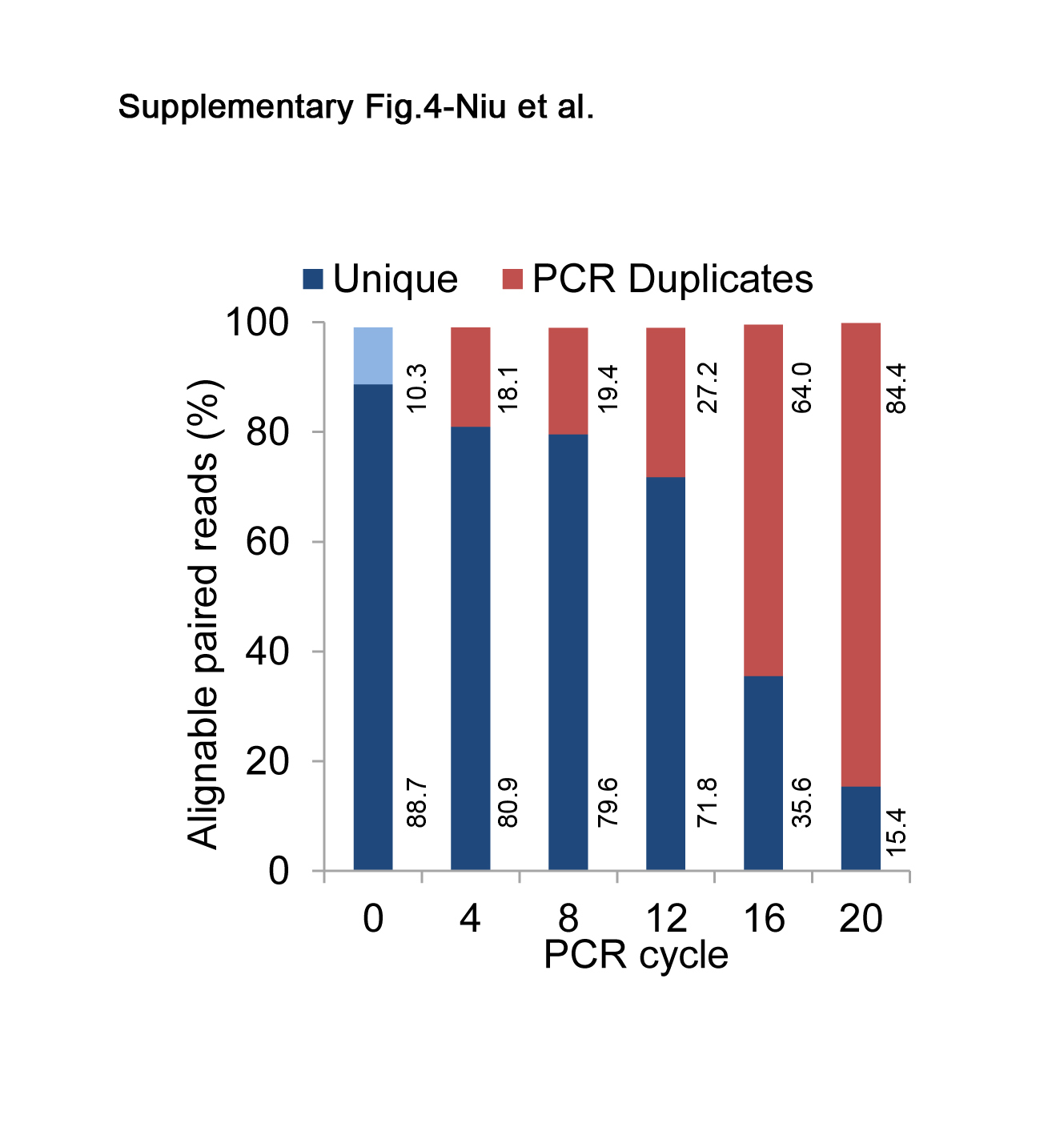

### Supplementary Figure 5

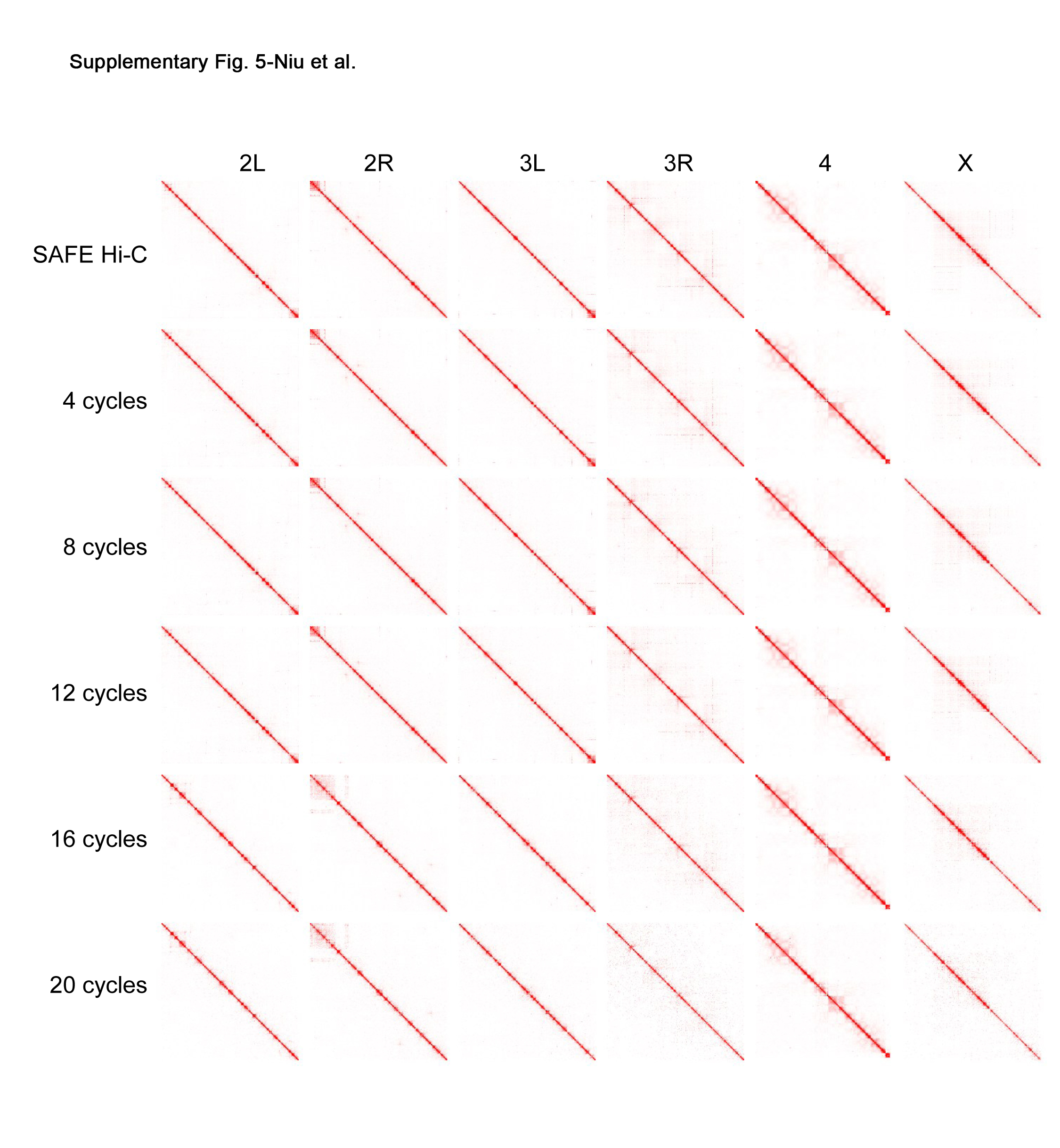

### Supplementary Figure 6

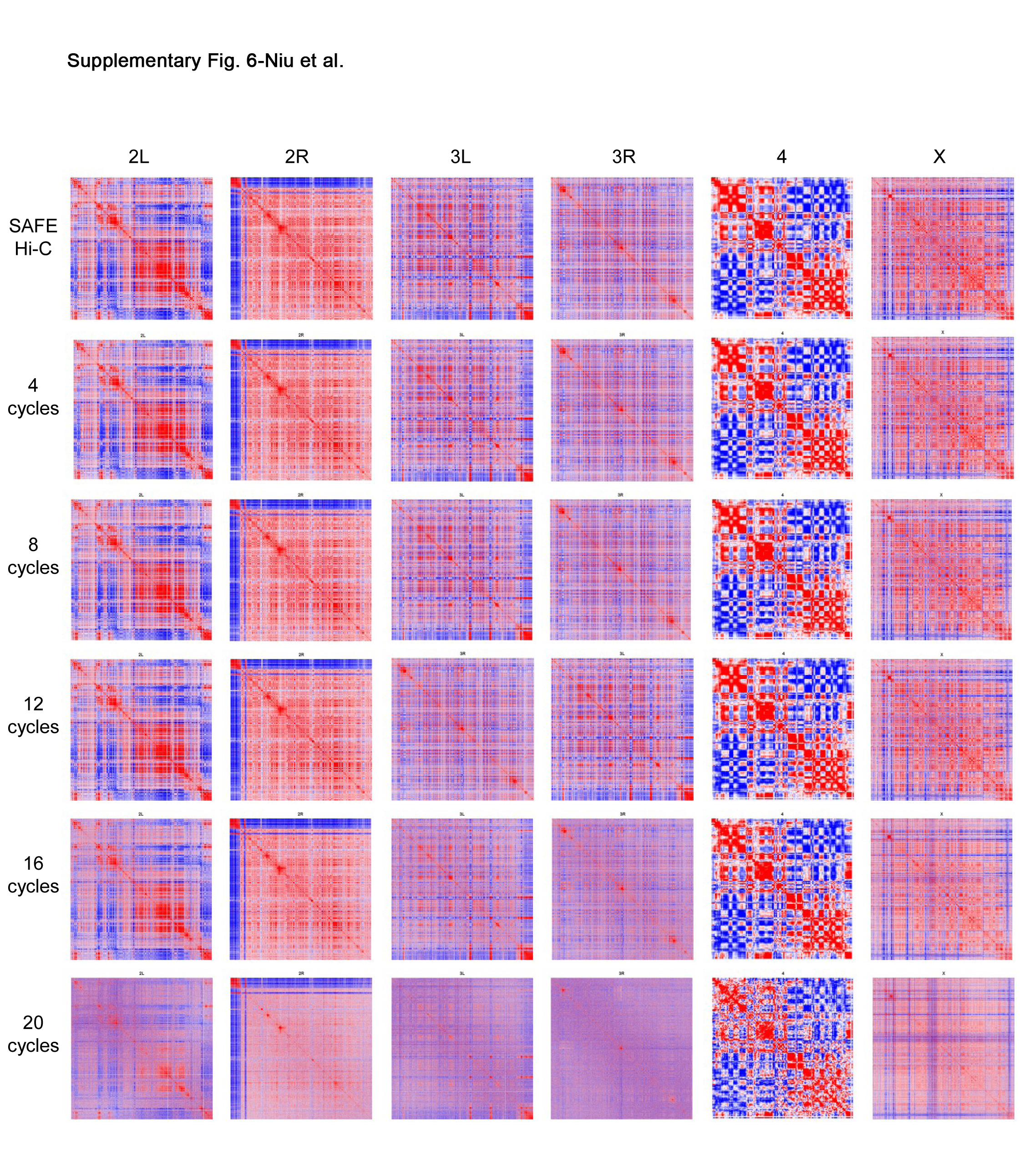

### Supplementary Figure 7

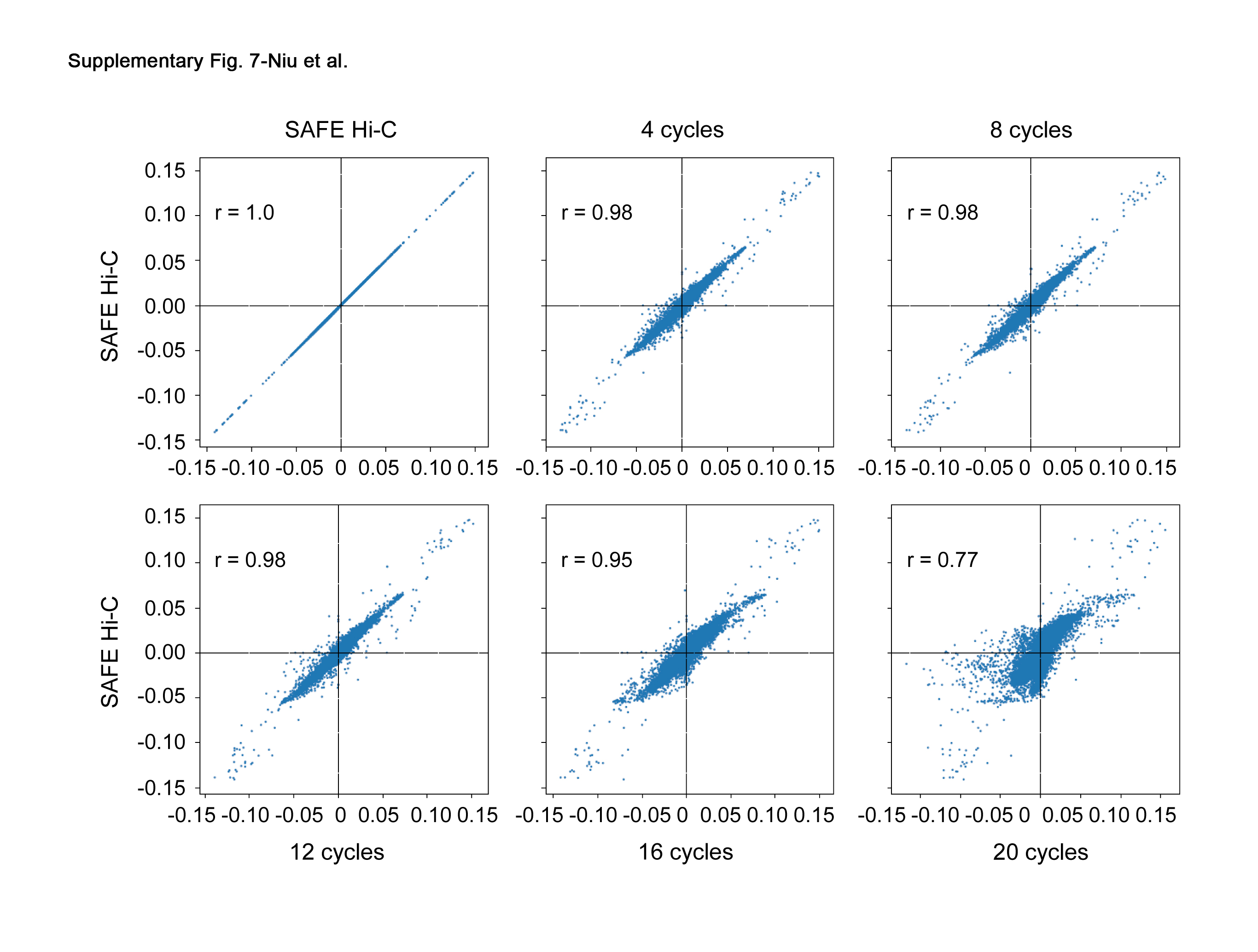

### Supplementary Figure 8

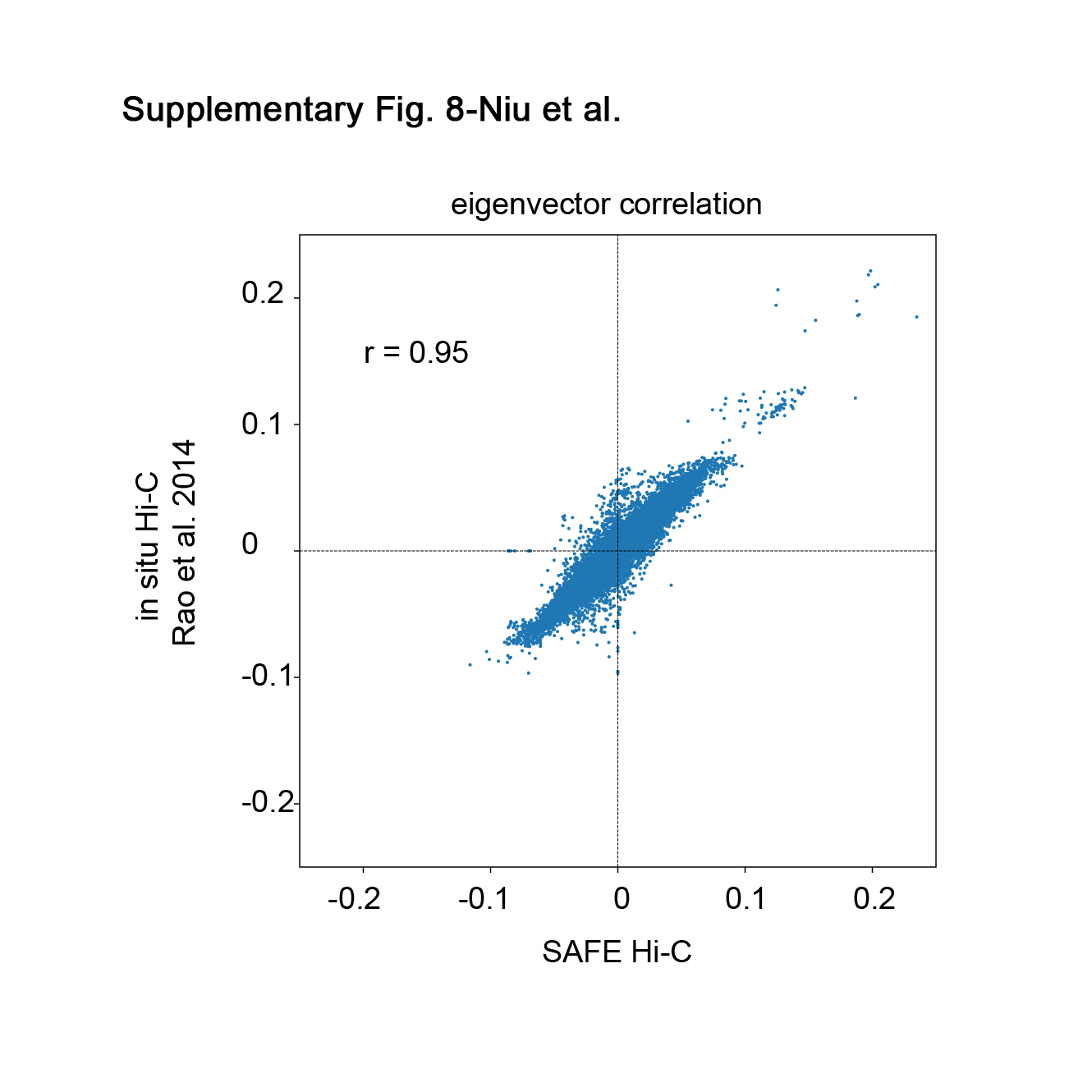

### Supplementary Figure 9

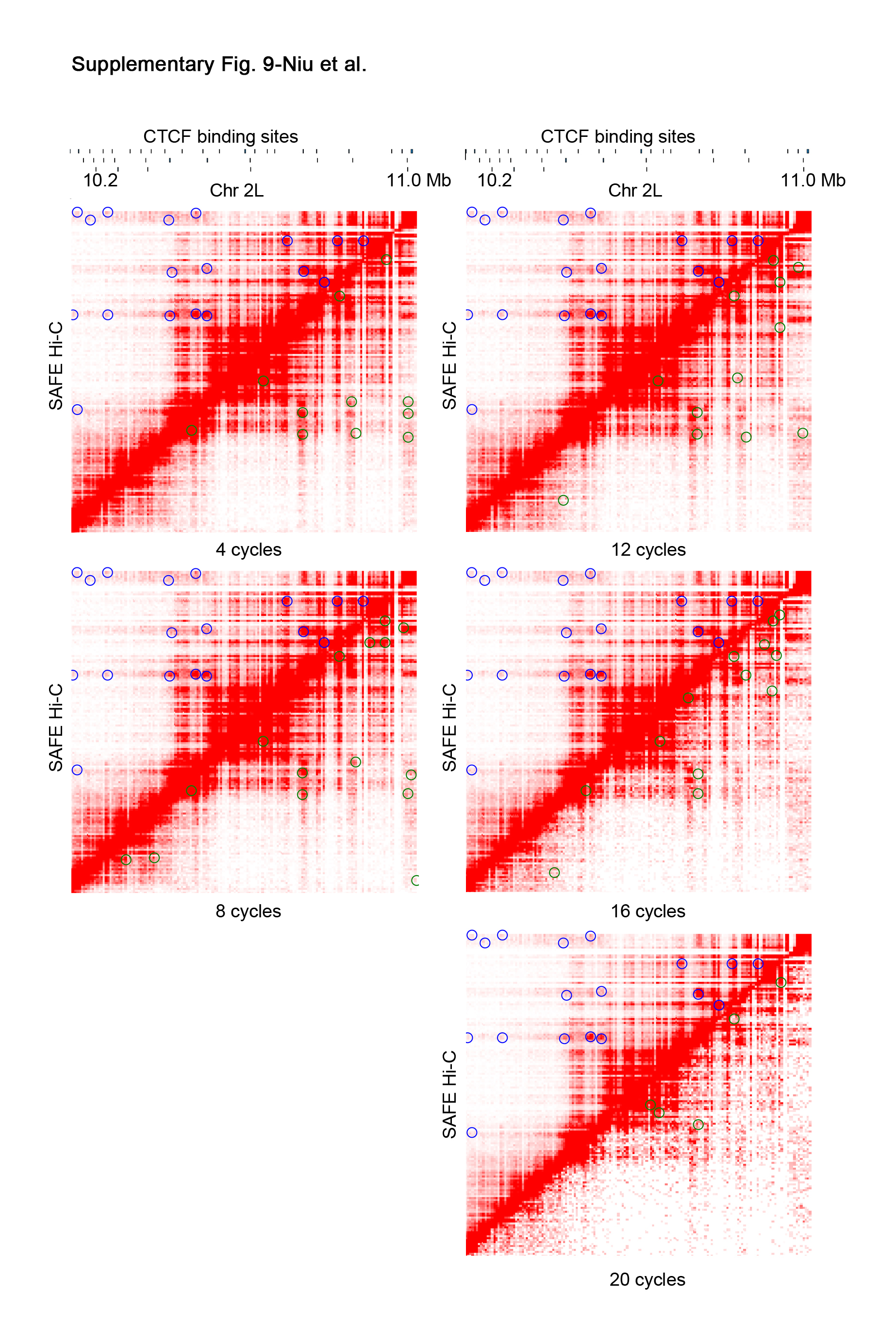

### Supplementary Figure 10

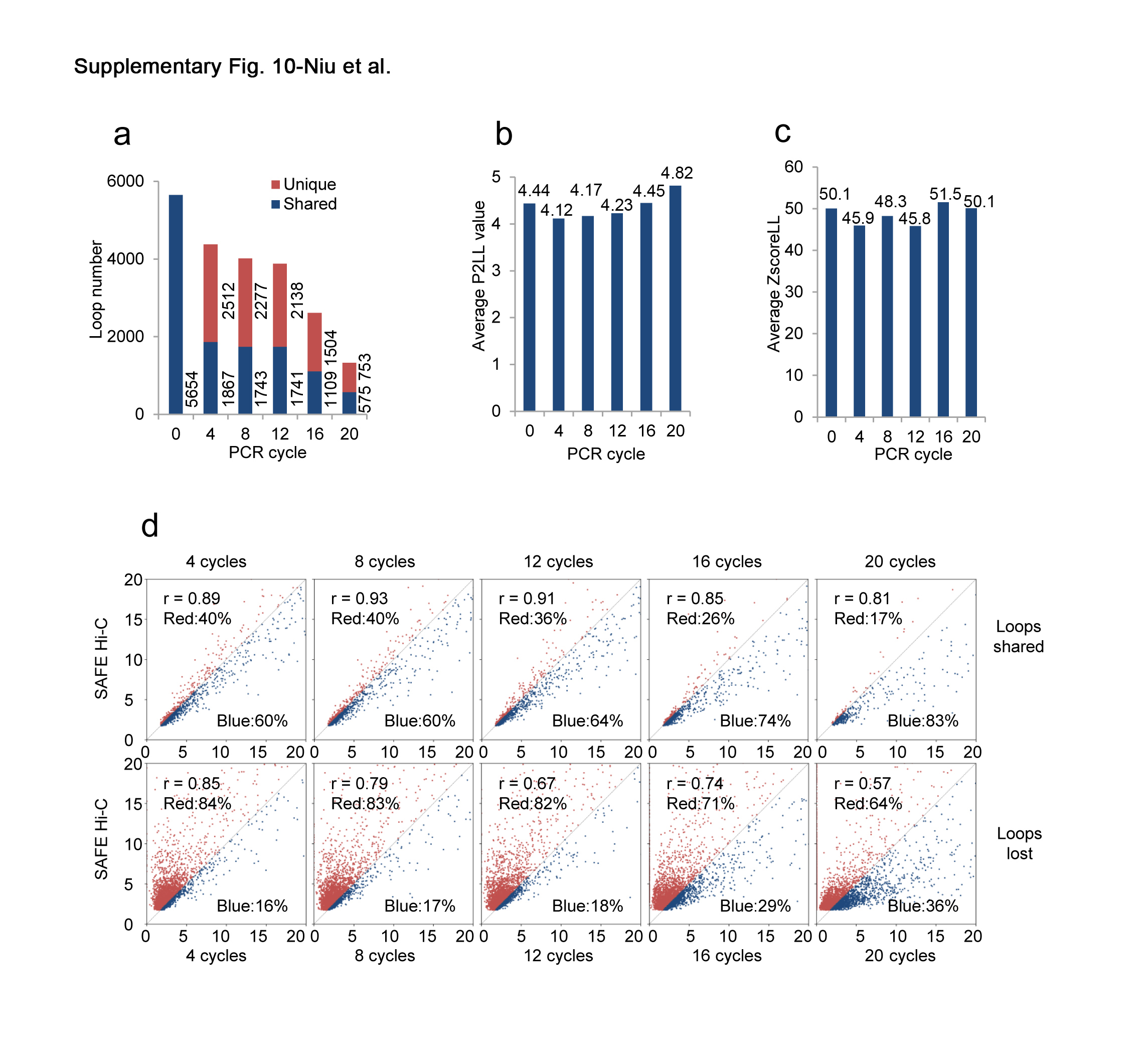

### Supplementary Table 1

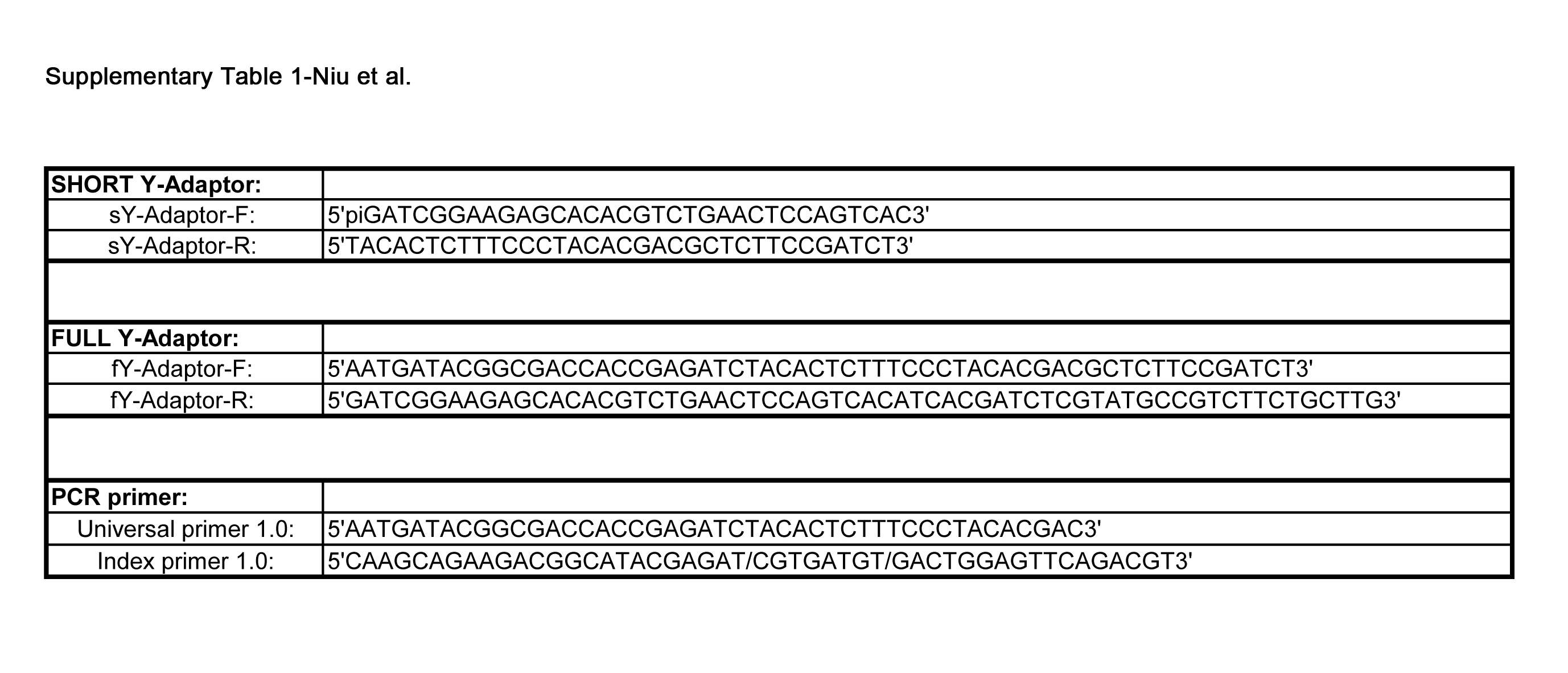

### Supplementary Table 2

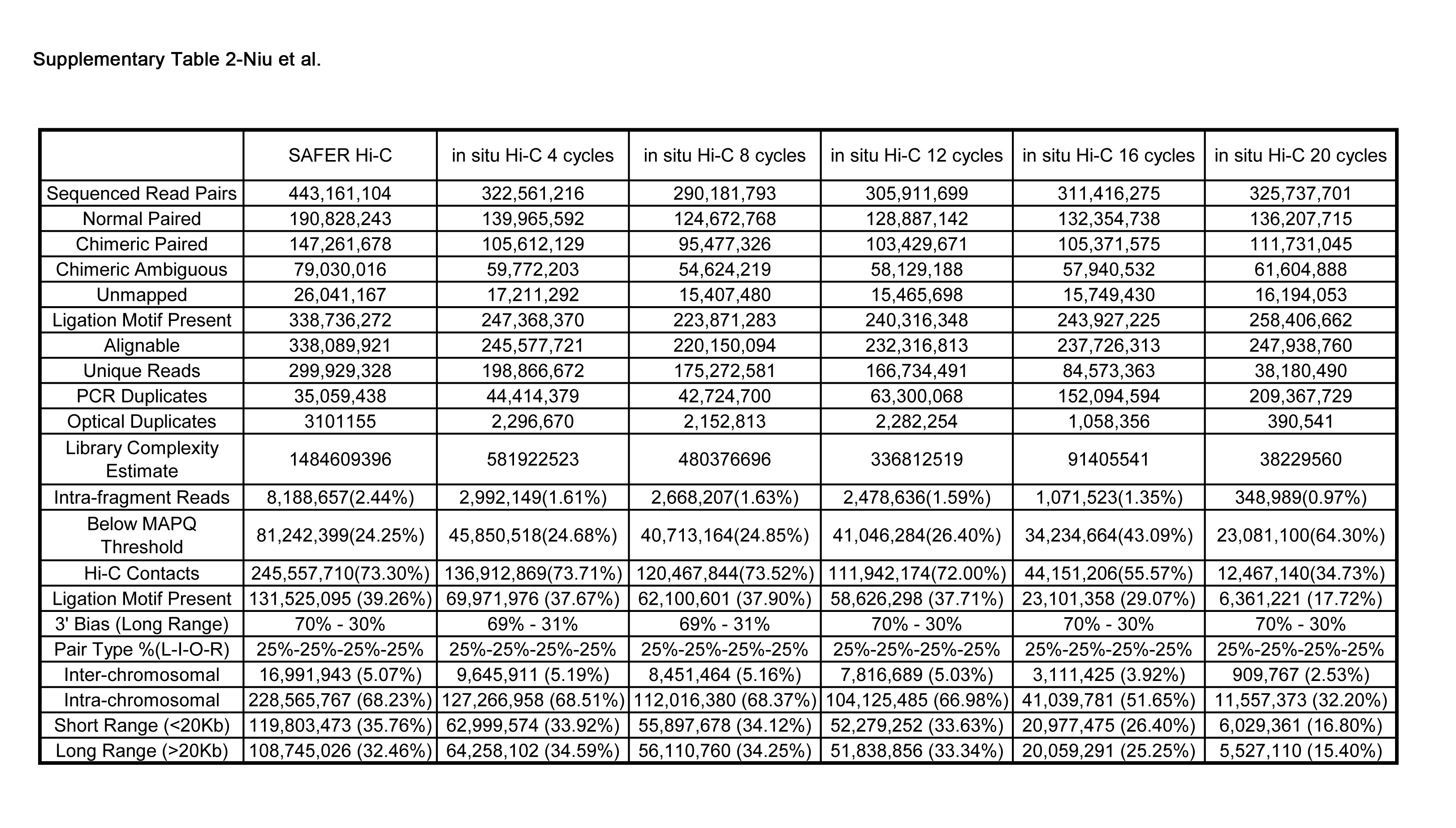

### Supplementary Table 3

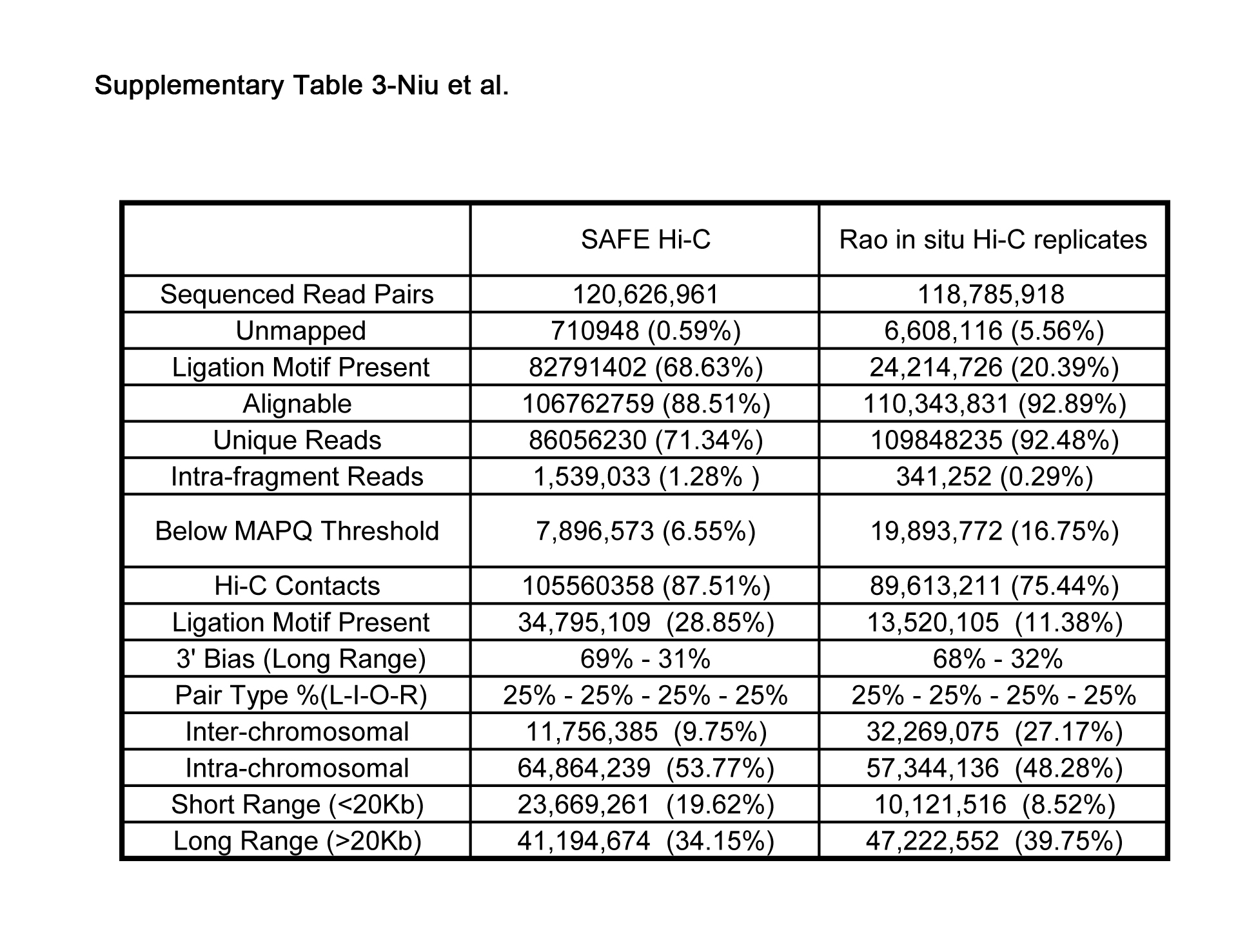
